# Food-borne mycotoxin hazards in the Kenyan market-a retrospective study

**DOI:** 10.1101/773747

**Authors:** James Karuku Kibugu, David Mburu, Leonard Karongo Munga, Richard Kurgat, Bernard Mukasa, Fransisca Naliaka Lusweti, Delia Grace, Johanna Lindahl

## Abstract

Mycotoxin contamination data (n=1818) in feed and food from major laboratories were categorized into hazardous and non-hazardous using contaminants regulatory limits, analyzed by logistic regression and chi-square test to identify potential health hazards. Feeds were most contaminated, with 64% and 39% having total aflatoxin (AFT) levels above Kenyan and American standards respectively. Peanuts, the most contaminated food, had 61% and 47% of samples failing Kenyan and American AFT standards respectively. By European standards, wheat had highest AFT contamination rate of 84%. Half of baby foods sampled had AFT level above Kenyan and European standards. Maize had failure rates of 20% (Kenyan standard), 14% (American standard) and 25% (European standard) for AFT. We observed high frequency of mycotoxins (AFT, aflatoxin M1, zearalenone, T-2 toxin, ochratoxin A, fumonisins, deoxynivalenol) and AFT hazards with significantly (*p*<0.001) higher failure rates in wheat, peanuts, mycotoxin hazards in dairy products in that order (European standard). Failure rates were significantly (*p*<0.001) higher in feed ingredients (*p*<0.01), baby foods (*p*<0.05), maize (*p*<0.001), fodder (*p*<0.05) for mycotoxins, and compound feeds, peanuts, wheat (*p*<0.001), feed ingredients, baby foods (*p*<0.01), maize (*p*<0.001), fodder (0.01), in that order, for AFT (American standard). Fail rates were significantly higher for mycotoxins in compound feeds, feed ingredients, peanuts, wheat, baby foods, maize (*p*<0.001), herbal health drink (*p*<0.01), and for AFT in compound feeds, feed ingredients, peanuts, wheat (*p*<0.001), baby foods (*p*<0.01), herbal health drink (*p*<0.05), maize (*p*<0.001) in that order (Kenyan standard). High frequency of mycotoxin and AFT hazards in maize, baby foods, herbal health drink and aflatoxin M1 in dairy products was noted. Detection by different laboratories varied significantly (*p*<0.001). Health and economic implications of this and limitations of current food safety standards are discussed. Humans and animals in Kenya are chronically exposed to mycotoxin hazards that require constant surveillance and strict regulation.

## Introduction

Mycotoxins are toxic secondary metabolites produced by toxigenic fungi that infest food and feed materials during pre- and post-harvest periods [1]. The most commonly encountered dietary mycotoxins with worldwide occurrence are aflatoxins, ochratoxins, zearalenone, fumonisins, trichothecenes and patulin produced by the fungal genera *Aspergillus*, *Penicillium* and *Fusarium* [1,2,3,4]. Children in Africa are continuously exposed to dietary mycotoxins [5]. Effects of chronic exposure include aggravation of disease pathogenesis in experimental animals and humans ([6,7,8], reduced animal productivity [9], and impaired animal nutrition [10]. Mycotoxins can also be teratogenic, carcinogenic, mutagenic, estrogenic, nephrotoxic, hepatotoxic and immunosuppressive in humans and animals [4, 8]. Aflatoxin is an important risk factor for primary hepatocellular carcinoma, and is also associated with childhood stunting and immune depression in humans [11,12,13,14]. There is widespread distribution of aflatoxigenic fungi in Kenya [3], and where indeed acute aflatoxicosis in human and animals resulting in deaths has occurred [15,16,17]. Other than threatening human and animal health, mycotoxins also affect international trading and contribute to food and feed insecurity [1]. Most mycotoxins are stable to normal cooking and processing. After consumption, some mycotoxin metabolites can be carried over *in utero* [18], in breast milk [9,19,20] and in animal products [13]; all contribute to mycotoxin exposure in humans.

In order to protect humans from exposure, Kenya as well as many other countries have regulatory limits for some mycotoxins [1,2,21,22]. However, standards are rarely enforced in the developing world [23]. There are commercial and government laboratories that offer quality assurance services for food material destined for export and local consumption. The purpose of this study was to review available data from testing laboratories in order to identify potential dietary mycotoxin hazards in Kenya, and estimate their frequency as health risk factors.

## Materials and methods

### Data collection and management

We developed a list of major mycotoxin-testing laboratories in Kenya [15] as a sampling frame, and collected data on total aflatoxin (AFT), aflatoxin M1 (AFM1), zearalenone (ZEA), T-2 toxin, ochratoxin A (OTA), total fumonisins (FUMS) and deoxynivalenol (DON) contamination in human foods and animal feeds from the laboratories (Lab1, Lab2 and Lab3). One of the laboratory was a public institution that deals with agricultural research while the other two were private laboratories (Lab2 and Lab3) that process samples for clients.

Results of 1818 samples (323 and 1495 samples of animal feeds and human food respectively) analysed for mycotoxin residue levels between 2010 and 2015 were acquired for further analysis. The data were first broadly grouped into animal feeds and human foods and then further categorized into compound feeds, feed ingredients, and fodder feeds for animal feeds, and baby foods, herbal health drink, maize, peanuts, dairy products, tea, wheat, on-the-plate (maize slurry, omena, vegetables) and other foods for humans. When the results were below the limit of detection, we did not consider the lower limit of detection (LLoD) to be the mycotoxin level since the actual value could be anywhere below the limit. Instead, half of the LLoD was taken as the mycotoxin level for the present analyses. When results were given as above the upper limit of detection, this upper limit was used as the value in the analyses. These assumptions did not introduce any bias in the dependent response, since mycotoxin regulatory limits are within the sensitive range of analytical methods i.e. between lower and upper LoD.

Kenyan national standards for animal feeds [21,24,25,26,27,28,29,30,31] and human foods [21,32,33,34,35,36,37,38,39,40,41,42,43,44,45,46] were applied to categorize the materials as either above or below the maximum admissible levels. Similarly, this was repeated with United States Food and Drugs Administration (FDA) and European Union (EU) standards [1,2,22,47,48,49,50,51,52,53]. Tables 1 and 2 show the maximum limits (MLs) of seven mycotoxin residues as stipulated in these standards for regulation of contaminants in food and feeds. A sample was considered a mycotoxin or a total aflatoxins hazard if its toxin level was above the legal limit of at least one of the seven mycotoxins referred to as mycotoxins or total aflatoxins hazard respectively.

**Table 1.** Feed safety regulatory standards used in the study

| Feed matrix | Toxin | Feed safety regulatory standard |  |  |  |  |  |
| --- | --- | --- | --- | --- | --- | --- | --- |
|  |  | KEBS standard |  | US-FDA standard |  | EU standard |  |
|  |  | ML (ppb) | Reference | ML (ppb) | Reference | ML (ppb) | Reference |
| Wheat Bran (Feed ingredient) | AFT | 20 | KEBS, 2004 | 20 | FDA, 2019; 2018; 2017; Alshannaq & Yu, 2017; Smith *et al.*, 2016; South, 2014 | NSA<sup>FT</sup> | - |
| Omena (Feed ingredient) | AFT | 20 | KEBS, 2011a | 20 |  |  |  |
| Cotton Seed/ cake (Feed ingredient) | AFT | 10 | KEBS, 2002 | 20 |  |  |  |
| Sunflower Cake (Feed ingredient) | AFT | 10 | KEBS, 2002 | 20 |  |  |  |
| Maize products (Feed ingredient) | AFT | 10 | KEBS 2012c, d; 2001 | 20 |  |  |  |
| Wheat Bran (Feed ingredient) | FUM | 2000 | KEBS, 2017 | NS |  |  |  |
| Poultry finished feed | AFT | 10 | KEBS, 2012a | 20 |  |  |  |
| Dog finished feed | AFT | 10 | KEBS, 2012b | 20 |  |  |  |
| Compound feed for dairy cattle | AFT | 10 | KEBS, 2009 | 20 |  |  |  |
| Compound feed for rabbits | AFT | 10 | KEBS 2012e | 20 |  |  |  |
| Compound feed | AFT | 10 | KEBS, 2017 | 20 |  |  |  |
| Fodder feed | AFT | 20 | KEBS, 2019 | 20 |  |  |  |
| Wheat Bran (Feed ingredient) | DON | 1000 | KEBS, 2017 | 5000 |  | 8000 | EU, 2006 |
| Wheat Bran (Feed ingredient) | OTA | 5000 | KEBS, 2017 | NS | - | 250 | EU, 2006 |
| Feed ingredient | ZEA | NS | - | NS | - | 2000 | EU, 2006 |

ML-Maximum level permitted; US-FDA- Food and Drug Administration of the United States of America; KEBS-Kenya Bureau of Standards; ppb-Parts per Billion; NS-not set; AFT-Total aflatoxins; DON-Deoxynivalenol; FUM-Total fumonisins; OTA-Ochratoxin A; ZEA-Zearalenone; NSA<sup>AFT</sup> –EU standard for animal feeds has limit for AFB<sub>1</sub> but none for AFT

**Table 2.**
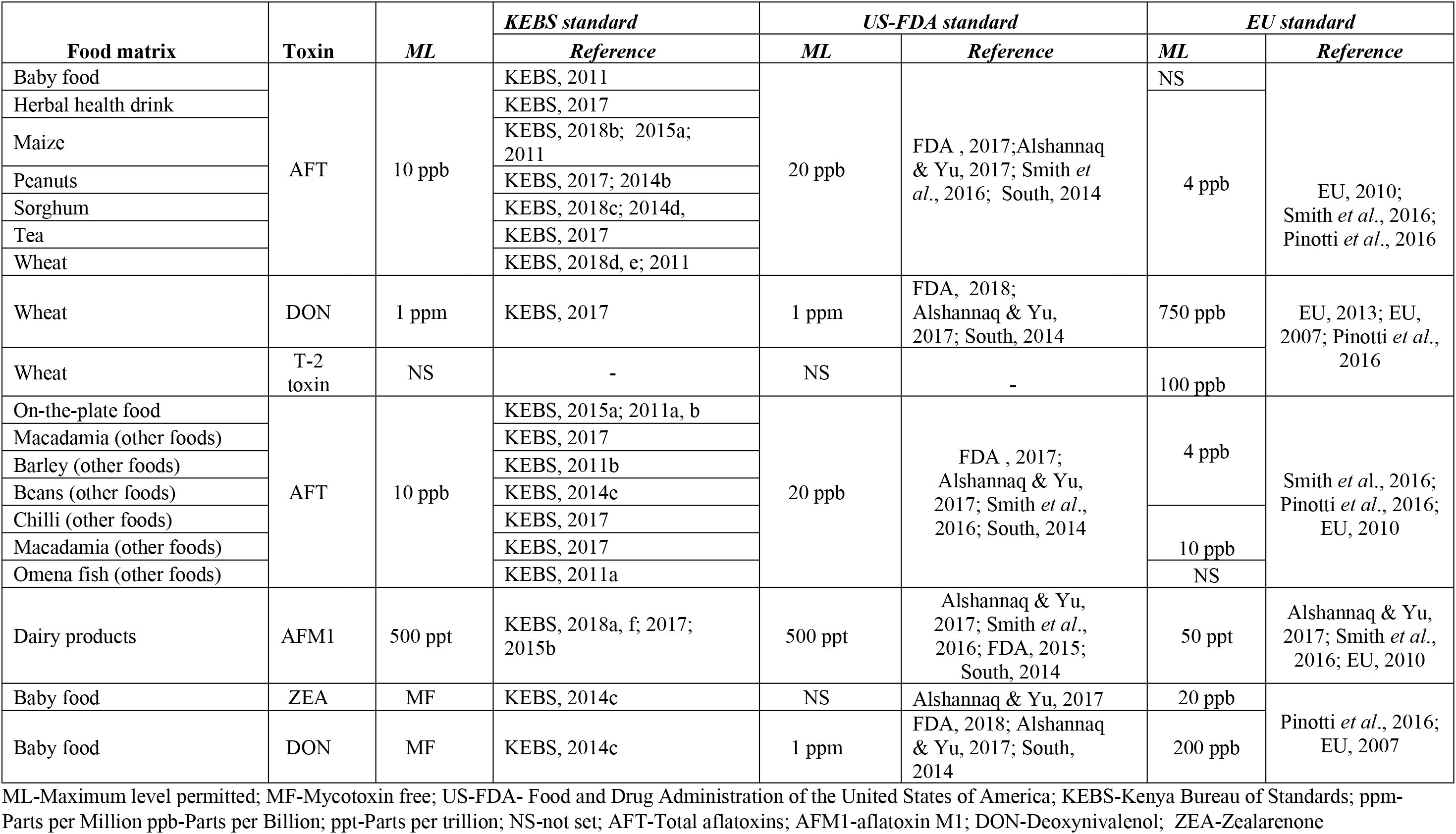
Food safety regulatory standards used in the study

### Statistical analysis

Logistic regression analysis was carried out to obtain odds ratios for the dependent response that is, above (1) or below (0) legal limits as stipulated in mycotoxin regulatory standards. The explanatory variables were 12 different feed/ food commodities and 3 mycotoxin-testing laboratories). The same was repeated for total aflatoxin data where the dichotomous dependent response variable. The mycotoxin and total aflatoxin contamination data had a binomial distribution (0 = not hazardous, 1 = hazardous), B(n, p), where n = number of feed or food samples and p = probability of attaining hazardous status. The following binary logistic regression model was fitted to both sets of data on statistical computer program (IBM SPSS Statistics 20):

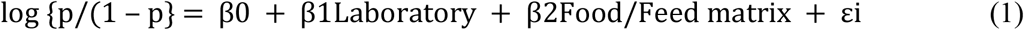

Where,

B_0_ = regression coefficient for Laboratory + Food/Feed matrix reference groups); β_1_Laboratory = regression coefficient for mycotoxin-testing Laboratories, β_2_Food/Feed matrix = regression coefficient for Food/Feed matrices and ε_i=_ random error.

Using the model, the effect of laboratory and feed/ food matrix on the dependent binary outcome variable response was determined. By default, classification cut off value of probability was set at 0.5, a threshold above and below which the vales are associated with hazardous state and non-hazardous state respectively. Further, the association between material matrices (animal feeds and human foods) and mycotoxin testing laboratories as predictors, and the binary response that is, presence or absence of mycotoxin hazard (either AFT, AFM1, ZEA, T-2 toxin, OTA, FUMS or DON) was further determined employing Pearson chi square-test of independence. This was repeated for total aflatoxins hazard.

## Results

### Frequency of mycotoxin hazards in various animal feed and human food matrices

The failure rates in attaining standards of four feed materials (n=323) and eight food materials (n=1495) tested for total aflatoxins and seven mycotoxins (collectively termed mycotoxins) are given in Table 3. Compound animal feeds were the most contaminated feed matrix with 64 % and 39 % of the 92 samples tested having total aflatoxin levels above the regulatory limit by Kenyan and American standards respectively. Peanuts were the most contaminated human food with 62 % and 47 % of the 180 tested samples having levels above the legal limit by Kenya and American standards respectively for total aflatoxin content. By European standard for total aflatoxin, wheat had the highest contamination rate with 84 % of the 105 samples tested having levels above the regulatory limit. Half (50%) of the baby foods failed to meet Kenyan and European standards for total aflatoxins. By Kenyan, American and European standards respectively, maize (a common staple food in Kenya) had total aflatoxin failure rates of 20, 14 and 25 % respectively. Again, maize was the most frequently tested food (Table 3). Failure rates of the feed and food samples for the seven different mycotoxins are shown in Table 4. Total aflatoxin was the most frequently tested contaminant followed by aflatoxin M1. In the few samples tested for ochratoxin A and zearalenone, 100% exceeded the regulatory limits by Kenya standard. Failure rate in dairy products was 60 % according to the European standard for aflatoxin M1 content (Table 4).

**Table 3.** Frequency of hazardous mycotoxin contamination in animal feeds and human foods

| Food/ feed matrix | Hazard | Proportion (%) of contaminated samples with levels above legal limit |  |  |  |  |  |  |  |  |  |  |  |
| --- | --- | --- | --- | --- | --- | --- | --- | --- | --- | --- | --- | --- | --- |
|  |  | Laboratory 1 |  |  | Laboratory 2 |  |  | Laboratory 3 |  |  | All Laboratories |  |  |
|  |  | KEBS | FDA | EU | KEBS | FDA | EU | KEBS | FDA | EU | KEBS | FDA | EU |
|  |  | nFE= 16; nFO=610 |  |  | nFE= 95; nFO=90 |  |  | nFE= 212; nFO= 795 |  |  |  |  |  |
| Feeds (n=323) | Mycotoxins | 5<sup>a</sup> | 4<sup>b</sup> | NS | 7<sup>a</sup> | 1<sup>b</sup> | NS | 18<sup>a</sup> | 15<sup>b</sup> | NS | 30*** | 20* | NS |
| Foods (n=1495) |  | 6<sup>a</sup> | 3<sup>b</sup> | 10<sup>c</sup> | 0.2<sup>a</sup> | 0.1<sup>b</sup> | 0.5<sup>c</sup> | 16<sup>a</sup> | 12<sup>b</sup> | 28<sup>c</sup> | 22*** | 15* | 39 |
|  |  | nFE=16 ; nFO=610 |  |  | nFE= 95; nFO=90 |  |  | nFE= 199; nFO=597 |  |  |  |  |  |
| Feeds (n=310) | Total aflatoxins | 5<sup>a</sup> | 4<sup>b</sup> | NS | 9<sup>a</sup> | 1<sup>b</sup> | NS | 18<sup>a</sup> | 15<sup>b</sup> | NS | 32** | 20 | NS |
| Foods (n=1297) |  | 7<sup>a</sup> | 4<sup>b</sup> | 12<sup>c</sup> | 0.2<sup>a</sup> | 0.1<sup>b</sup> | 0.6<sup>c</sup> | 18<sup>a</sup> | 14<sup>b</sup> | 23<sup>c</sup> | 25** | 18 | 36 |
|  |  | nFI= 1; nCF=15 |  |  | nFI=57 ; nCF=;38 |  |  | nFI= 6; nCF=39;<br>nFOD=154 |  |  |  |  |  |
| Feed ingredients (n=64) | Total aflatoxins | 2 | 2 | NS | 20 | 2 | NS | 5 | 3 | NS | 27 | 7 | NS |
| Compound feed (n=92) |  | 16 | 12 | NS | 16 | 2 | NS | 32 | 25 | NS | 64 | 39 | NS |
| Fodder feed (n=154) |  | - | - | NS | - | - | NS | 16 | 15 | NS | 16 | 15 | NS |
|  | Total aflatoxins | nM=241; nPF= 369 |  |  | nM=;86 nO= 4 |  |  | nB=6; nH=21; nM=234;<br>nP=180 ; nT=37 ;<br>nW=105; nO=14 |  |  |  |  |  |
| Baby foods (n=6) |  | - | - | - | - | - | - | 50 | 33 | NS | 50 | 33 | NS |
| Herbal health drink (n=21) |  | - | - | - | - | - | - | 24 | 10 | 48 | 24 | 10 | 48 |
| Maize (n=561) |  | 9 | 6 | 12 | 0.5 | 0.2 | 1 | 10 | 7 | 12 | 20 | 13 | 25 |
| Peanuts (n=180) |  | - | - | - | - | - | - | 62 | 47 | 70 | 62 | 47 | 70 |
| Tea (n=37) |  | - | - | - | - | - | - | 0 | 0 | 0 | 0 | 0 | 0 |
| Wheat (n=105) |  | - | - | - | - | - | - | 54 | 46 | 85 | 54 | 46 | 85 |
| On-the- plate food (n=369) |  | 9 | 4 | 25 | - | - | - | - | - | - | 9 | 4 | 25 |
| Other foods (n=14) |  | - | - | - | 0 | 0 | 0 | 17 | 0 | 25 | 17 | 0 | 25 |

NS=Regulatory limit not set; n<sub>FE</sub>, n<sub>FO</sub>, n<sub>FI</sub>, n<sub>CF</sub>, n<sub>FOD</sub>, n<sub>B</sub>, n<sub>H</sub>, n<sub>M</sub>, ; n<sub>P</sub>, ; n<sub>T</sub>, n<sub>W</sub>, n<sub>PF</sub>= and n<sub>O</sub>=stand for sample sizes of feed, food, feed ingredient, compound feed, fodder feed, baby food, herbal health drink, maize, peanut, wheat, on-the- plate food and other food respectively. <sup>a</sup>, <sup>b</sup>, and <sup>c</sup> indicate significant association between laboratory and hazard status at $p=0.001$ , whereas \*, \*\* and \*\*\*represent significant association between matrix and hazard status respectively at $p=0.05$ , 0.01 and 0.001.

**Table 4.** Mycotoxins detected in foods and feeds in Kenya (2010-2015)

| *Type of contaminant* | **Proportion (%) of contaminated samples** |  |  |
| --- | --- | --- | --- |
|  | *KEBS standard* | *FDA standard* | *EU standard* |
| *Total aflatoxin (n=1610)* | 26.29 | 18.14 | 36.02 |
| *Aflatoxin M1 (n=192)* | 0.52 | 0.52 | 59.9 |
| *Deoxynivalenol (n=6)* | 0 | 0 | 25 |
| *Total fumonisins (n=6)* | 0 | 0 | 0 |
| *Ochratoxin A (n=1)* | 100* | 0 | 0 |
| *T-2 toxin (n=2)* | 0 | 0 | 0 |
| *Zearalenone (n=4)* | 100* | 0 | 25 |

\*Sample size is insufficient

### Comparison of mycotoxin hazard rates in animal feed and human food

Failure rate to attain standards in animal feeds and human foods by mycotoxins and total aflatoxins (AFT) is shown in Table 3. Chi square test of independence showed significant association between both type of matrix and mycotoxin analysing laboratories, and hazards status. By Kenyan standard, animal feeds had higher mycotoxin failure rate (30 %) compared to 22 % in human food (*p* <0.001, Pearson chi square = 16.056, DoF =2) for mycotoxins, and 32% in animal feeds compared to 25% in human food (*p* <0.01, Pearson chi square = 7.498, DoF = 2) for total aflatoxins. By American standard, animal feeds had higher mycotoxin failure rate (20 %) compared to 15 % in human foods (*p* <0.05, Pearson chi square = 4.328, DoF = 2) but no significant association was observed for total aflatoxin hazard. There was significant (*p* <0.001, DoF = 2) association between the various analysing laboratories and hazards status by Kenyan (Pearson chi square = 41.64), American (Pearson chi square =79.338), and European (Pearson chi square = 141.913) standards for mycotoxins and Pearson chi square = 82.812, 119.092 and 114.23 respectively for total aflatoxins.

### Effect of feed and food matrices on frequency of mycotoxin hazards

Tables 5-7 show logistic regression results of the two explanatory variables giving odds ratios (OR) of dietary mycotoxin and total aflatoxin hazards occurrence in the matrices and their detection capability as a function of European (EU), American (FDA) and Kenyan (KEBS) standards for regulation of food contaminants. Results of Hosner & Lemeshow test (*p* >0.01) indicate good fitting models. Relative to a reference food matrix, high frequency of mycotoxin and total aflatoxin hazards were observed in several food/ feed materials as a function of maximum acceptable contamination limits set by Kenyan, American and European regulatory organizations. By European standard (EU), odds of presence of mycotoxin and total aflatoxin hazards were higher in wheat (OR =11.7) and (OR = 14.6) respectively, peanuts for both hazards (OR = 6.1), and mycotoxin hazard in dairy products (OR = 3.9) compared to on-the-plate food (Table 5).

**Table 5.** Logistic regression results of risks associated with dietary mycotoxins as stipulated in European Union standards for regulation of contaminants foods and feeds

| *Mycotoxin* | **Mycotoxin-testing laboratories and Food/ feed materials** | **Regression coefficient (B ± S.E)** | **Odds Ratio** | **95% CI Odds Ratio** |
| --- | --- | --- | --- | --- |
| ***Any one mycotoxin (AFT, AFM1, ZEA, T-2 toxin, OTA, FUMS, DON)*** | Lab1 (610) | -0,15 ± 0.20 | 0.86 | 0.58- 1.28 |
|  | Lab2 (90) | -1.49 ± 0.40*** | 0.23 | 0.1- 0.49 |
|  | Baby foods (8) | 0.45 ± 0.77 | 1.57 | 0.35- 7.08 |
|  | Dairy products (192) | 1.37 ± 0.28*** | 3.92 | 2.27- 6.76 |
|  | Herbal health drink (21) | 0.87 ± 0.50 | 2.38 | 0.9- 6.31 |
|  | Maize (561) | 0.12 ± 0.19 | 1.13 | 0.78- 1.63 |
|  | Peanuts (180) | 1.81 ± 0.29** | 6.12 | 3.49-10.74 |
|  | Tea (37) | -20.24 ± 6607.68 | 1.62x10<sup>-9</sup> | - |
|  | Wheat (109) | 2.46 ± 0.34*** | 11.67 | 5.97-22.82 |
|  | Other foods (12) | 0.24 ± 0.73 | 1.27 | 0.31- 5.26 |
|  | *Reference: Lab 3 (789)/ on-the-plate (369)* | -0.96 ± 0.24*** | 0.38 |  |
| ***Total aflatoxins*** | Lab1 (610) | -0.15 ± 0.20 | 0.86 | 0.58- 1.28 |
|  | Lab2 (90) | -1.49 ± 0.40*** | 0.23 | 0.10-0.49 |
|  | Baby foods (6) | 0.96 ± 0.85 | 2.62 | 0.50-13.87 |
|  | Herbal health drink (21) | 0.87 ± 0.50 | 2.38 | 0.90-6.31 |
|  | Maize (561) | 0.12 ± 0.19 | 1.13 | 0.78-1.63 |
|  | Peanuts (180) | 1.81 ± 0.29*** | 6.12 | 3.49-10.74 |
|  | Tea (37) | -20.24 ± 6607.68 | 1.62x10<sup>-9</sup> | - |
|  | Wheat (105) | 2.68 ± 0.36*** | 14.59 | 7.20-29.54 |
|  | Other foods (12) | 0.24 ± 0.73 | 1.27 | 0.31-5.26 |
|  | *Reference: Lab 3 (591)/ on-the-plate (369)* | -0.96 ± 0.24*** | 0.38 |  |

\*p<0.05; \*\* p<0.01; \*\*\* p<0.001; NS=p-value not significant; AFT=Total aflatoxins; AFM1=Aflatoxin M1; ZEA=Zearalenone; OTA=Ochratoxin A; FUMS=Total fumonisins; DON=Deoxynivalenol

**Table 6.** Logistic regression results of risks associated with dietary mycotoxins as stipulated in Food & Drug administration of United States of America (FDA) standards for regulation of contaminants in foods and feeds

| *Mycotoxin* | **Mycotoxin-testing laboratories and Food/ feed materials** | **Regression coefficient (B ± S.E)** | **Odds Ratio** | **95% CI Odds Ratio** |
| --- | --- | --- | --- | --- |
| ***Any one mycotoxin (AFT, AFM1, ZEA, T-2 toxin, OTA, FUMS, DON)*** | Lab1 (626) | -0.14 ± 0.23 | 0.87 | 0.55-1.36 |
|  | Lab2 (180) | -3.20 ± 0.57*** | 0.04 | 0.01-0.12 |
|  | Baby foods (7) | 2.03 ± 0.90* | 7.65 | 1.30-45.06 |
|  | Dairy products (192) | -2.30 ± 1.06* | 0.1 | 0.01-0.80 |
|  | Feed Ingredients (59) | 2.12 ± 0.75** | 8.31 | 1.89-36.45 |
|  | Compound feeds (93) | 3.44 ± 0.41*** | 31.26 | 14.1-69.34 |
|  | Fodder feeds (154) | 1.21 ± 0.41** | 3.36 | 1.50-7.53 |
|  | Herbal health drink (21) | 0.70 ± 0.82 | 2.01 | 0.40-10.03 |
|  | Maize (561) | 1.35 ± 0.31*** | 3.89 | 2.11-7.08 |
|  | Peanuts (180) | 2.84 ± 0.38*** | 17.1 | 8.19-35.73 |
|  | Tea (37) | -18.25 ± 6607.68 | 1.18x10<sup>-10</sup> | - |
|  | Wheat (107) | 2.74 ± 0.40*** | 15.55 | 7.16-33.79 |
|  | Other foods (18) | -17.92 ± 9047.27 | 1.65x10<sup>-8</sup> | - |
|  | *Reference: Lab 3 (992)/ on-the-plate (369)* | -2.95 ± 0.34*** | 0.05 |  |
| ***Total aflatoxins*** | Lab1(626) | -0.16 ± 0.23 | 0.86 | 0.54- 1.35 |
|  | Lab2 (180) | -3.27 ± 0.57*** | 0.04 | 0.01- 0.12 |
|  | Baby foods (6) | 2.24 ± 0.93** | 9.43 | 1.52-58.64 |
|  | Feed Ingredients (58) | 2.27 ± 0.77** | 9.7 | 2.16-43.65 |
|  | Compound feeds (92) | 3.48 ± 0.41*** | 32.62 | 14.64-72.66 |
|  | Fodder feeds (154) | 1.20 ± 0.41** | 3.31 | 1.48- 7.44 |
|  | Herbal health drink (21) | 0.69 ± 0.82 | 1.99 | 0.40- 9.90 |
|  | Maize (561) | 1.35 ± 0.31*** | 3.84 | 2.10- 7.04 |
|  | Peanuts (180) | 2.83 ± 0.38*** | 16.88 | 8.08-35.27 |
|  | Tea (37) | -18.27 ± 6607.68 | 1.17x10<sup>-8</sup> | - |
|  | Wheat (105) | 2.77 ± 0.40*** | 15.89 | 7.30-34.58 |
|  | Other foods (18) | -17.93 ± 9034.86 | 1.63x10<sup>-8</sup> | - |
|  | *Reference: Lab 3(795) + on-the-plate (369)* | -2.94 ± 0.35*** | 0.05 |  |

\*p<0.05; \*\* p<0.01; \*\*\* p<0.001; NS=p-value not significant; AFT=Total aflatoxins; AFM1=Aflatoxin M1; ZEA=Zearalenone; OTA=Ochratoxin A; FUMS=Total fumonisins; DON=Deoxynivalenol

**Table 7.** Logistic regression results of risks associated with dietary mycotoxins as stipulated in Kenya (KEBS) standards for regulation of contaminants in foods and feeds

| *Mycotoxin* | **Mycotoxin-testing laboratories and Food/ feed materials** | **Regression coefficient (B ± S.E)** | **Odds Ratio** | **95% CI Odds Ratio** |
| --- | --- | --- | --- | --- |
| ***Any one mycotoxin (AFT, AFM1, ZEA, T-2 toxin, OTA, FUMS, DON)*** | Lab1 (626) | 0.06 ± 0.21 | 1.06 | 0.71-1.61 |
|  | Lab2 (185) | -1.82 ± 0.33*** | 0.16 | 0.08-0.31 |
|  | Baby foods (8) | 2.35 ± 0.76*** | 10.49 | 2.37-46.44 |
|  | Dairy products (192) | -2.90 ± 1.04** | 0.05 | 0.01-0.42 |
|  | Feed Ingredients (66) | 2.88 ± 0.46*** | 17.9 | 7.29-43.93 |
|  | Compound feeds (93) | 3.73 ± 0.37*** | 41.82 | 20.22-86.46 |
|  | Fodder feeds (154) | 0.66 ± 0.35 | 1.94 | 0.97-3.88 |
|  | Herbal health drink (21) | 1.19 ± 0.58* | 3.28 | 1.05-10.26 |
|  | Maize (561) | 1.07 ± 0.23*** | 2.93 | 1.85-4.63 |
|  | Peanuts (180) | 2.83 ± 0.32*** | 16.87 | 9.08-31.34 |
|  | Tea (37) | -18.85 ± 6607.68 | 6.49x10<sup>-9</sup> | - |
|  | Wheat (107) | 2.48 ± 0.34*** | 11.96 | 6.17-23.17 |
|  | Other foods (18) | 0.50 ± 0.81 | 1.65 | 0.34-8.00 |
|  | *Reference: Lab 3(995)+ on-the-plate (369)* | -2.35 ± 0.28*** | 0.10 |  |
| ***Total aflatoxins*** | Lab1 (626) | 0.03 ± 0.21 | 1.03 | 0.68-1.56 |
|  | Lab2. (180) | -2.05 ± 0.36*** | 0.13 | 0.06-0.26 |
|  | Baby foods (6) | 2.32 ± 0.86** | 10.14 | 1.87-54.95 |
|  | Feed Ingredients (62) | 3.19 ± 0.49*** | 24.29 | 9.24-63.85 |
|  | Compound feeds (92) | 3.87 ± 0.39*** | 47.82 | 22.44-101.9 |
|  | Fodder feeds (154) | 0.63 ± 0.36 | 1.87 | 0.93-3.76 |
|  | Herbal health drink (21) | 1.15 ± 0.58* | 3.17 | 1.01-9.92 |
|  | Maize (561) | 1.07 ± 0.23*** | 2.9 | 1.83-4.60 |
|  | Peanuts (180) | 2.79 ± 0.32*** | 16.31 | 8.77-30.34 |
|  | Tea (37) | -18.87 ± 6607.68 | 62.8 x10<sup>-6</sup> | - |
|  | Wheat (105) | 2.49 ± 0.34*** | 12.04 | 6.19-23.42 |
|  | Other foods (18) | 0.48 ± 0.81 | 1.61 | 0.33-7.84 |
|  | *Reference: Lab 3(794)+ on-the-plate (369)* | -2.32 ± 0.28*** | 0.10 |  |

\*p<0.05; \*\* p<0.01; \*\*\* p<0.001; NS=p-value not significant; AFT=Total aflatoxins; AFM1=Aflatoxin M1; ZEA=Zearalenone; OTA=Ochratoxin A; FUMS=Total fumonisins; DON=Deoxynivalenol

By the American standard (FDA), the odds of detecting mycotoxin and total aflatoxin hazards were higher in compound feeds (OR = 31.3, 32.6 respectively), and fodder feeds (OR = 8.3, 9.7 respectively) compared to on-the-plate food. In human foods, the odds of presence of mycotoxin and total aflatoxin hazards were higher in peanuts (OR = 17.1, 16.9 respectively), wheat (OR = 18.0, 15.9), maize (OR = 3.9, 3.8 respectively) and baby foods (OR = 7.7, 9.4 respectively) compared to on-the-plate food (Table 6). The odds of presence of mycotoxin hazard were lower in dairy products (OR = 0.1) compared to on-the-plate food.

For the Kenya standard (KEBS), odds of presence of mycotoxin and total aflatoxin hazards were higher in feed ingredients (OR = 17.9, 24.3 respectively), in compound feeds (OR = 41.8, 47.8 respectively), peanuts (OR = 16.9, 16.3 respectively), wheat (OR = 12.0), baby foods (OR = 10.5, 10.1 respectively) and maize (OR = 2.9) compared to on-the-plate food. Odds of herbal health drink are more prone to the two hazards (OR = 3.3, 3.2 respectively), while odds of presence of mycotoxin hazard were lower in dairy products (OR = 0.05) compared to on-the-plate food (Table 7).

### Hazard detection capacity by mycotoxin-testing laboratories

Further evidence suggests difference in hazard detection ability by different mycotoxin-testing facilities. Odds of detection by Lab2 were lower for mycotoxin and total aflatoxin hazards by EU (OR = 0.2), FDA (OR = 0.04) and KEBS (OR = 0.2, 0.1 respectively) standards compared to reference laboratory (Lab3) (Tables 5 and 6). No difference was observed between Lab1 and Lab3 (Tables 5-7).

## Discussion

We used USA and European Union food contaminants regulatory systems in the present study because of their robust and strict nature, and regular interaction with the Kenyan economy. High frequency of mycotoxin hazards in food and feed materials with highest frequency in compound feeds, feed ingredients, peanuts and wheat in that order was observed. This agrees with other authors’ observation that animals are frequently fed more contaminated products compared to humans [54,55,56]. In resource poor environments, biomass considered unfit for human consumption is commonly used for feeding animals to avoid waste. Presence of mycotoxin hazards in animal feeds and foods is of great concern since chronic dietary mycotoxicosis is associated with adverse health effects including immune dysfunction, nutritional deficiency, reduced fertility, drug and vaccine failure, and pathological and metabolic aberrations [4,7,57,58,59,60,61]. Mycotoxins impair productivity in farm animals [9,56,62] resulting in economic loss [1,63]. They carry-over to animal edible products [13], and are an impediment to international trade [1,2,64] since there is increasing demand for safe foods [14].

We identified peanuts and wheat as the human foods with significant frequency of mycotoxin and total aflatoxin hazards. High frequency of total aflatoxin hazard in maize and baby foods, herbal health drinks and aflatoxin M1, a hydroxylated metabolite of aflatoxin B1, in dairy products was also observed. These are readily available processed, raw and staple food items commonly consumed by both children and adults. In Africa and more so in Kenya, many foods such as cereals and dairy products although not generally classified as baby foods, are actually consumed by infants and young children. Some of the mycotoxins encountered in the present study for instance aflatoxins, ochratoxin A and fumonisins are teratogenic, carcinogenic, mutagenic, nephrotoxic, hepatotoxic and immunosuppressive in humans and animals [1,4,8,17,65,66]. Aflatoxin is anti-nutritional [10], and could be associated with childhood stunting and [11,12,13,14] in humans. Besides exerting direct toxicological effects, mycotoxins are also known to induce oncogenesis in synergy with human pathogens. The most abundant of the observed hazards, aflatoxins could have serious implication in aetiology and epidemiology of malignancies in human population. Its involvement in development of primary hepatocellular carcinoma through synergy with hepatitis B virus is well documented [67,68], and interaction with human papillomavirus in induction of oesophageal malignancy postulated [54,69]. Mycotoxins are known to modify disease through aggravation of pathogenesis of viral, bacterial, and parasitic infections in humans and experimental animals [6,70,71]. Of special interest is exacerbation of mucosa-associated diseases in the gastrointestinal and respiratory tracts by mycotoxins [70].

Cancer is an increasingly emerging health scourge in Sub-Saharan Africa [72,73] and major cause of death, ranking third with 28500 deaths and increased incidence reaching 37000 new cases during 2012 in Kenya [54,74,75] and more alarmingly in relatively young people [76]. In Kenya, the mean age of easophageal malignancy patients is approximately 50 years with high numbers of patients below 40 years. Our data show high frequency of mycotoxin hazards in foods destined for consumption by paediatric and pregnant individuals suggesting that exposure to the dietary carcinogenic hazards in Kenyan human population commences early in life. In fact, *in utero* mycotoxin exposure is not uncommon in Sub-Saharan Africa [18]. Since malignancy depends on exposure in terms of dose and time [77], with the young being more susceptible to environmental carcinogens [78,79] and more so to mycotoxins [59,80], our observation points to a possible important risk factor contributing to occurrence of cancer in relatively young individuals and human population as a whole in Kenya.

Oncogenesis can be induced via reduced immuno-surveillance [81] or potentiation of carcinogenic infections [70]. Several viral, bacterial and parasitic pathogens classified and established as human carcinogens are associated with some prevalent malignancies in Kenya [82] such as esophageal, stomach, liver, prostate, breast, cervical, ovarian cancers, chronic leukemia, endometrial, laryngeal, colorectal, nasopharyngeal, Kaposi’s sarcoma and non-Hodgkin’s lymphoma cancers [54,75,83,84,85,86]. High frequency of mycotoxins observed in the present study, could likely be an important risk factor in development of these infection-associated malignancies through either synergy, exacerbation of the carcinogenic infections or immunosuppression. Further studies are required to elucidate the relationship between the carcinogenic biological agents and chronic exposure to dietary mycotoxin hazards. Exposure to mycotoxins in children at a young age has been reported before in Kenya [19] and elsewhere [5,20] and our results should be a wakeup call to the relevant authorities to protect vulnerable individuals from these lethal toxins.

The present study was not designed to assess laboratories’ ability to detect mycotoxins. We however observed significant difference in probability of failure for mycotoxin regulatory limits reported by the three laboratories. This could be explained by differences in analytical methods applied by the laboratories, differences in primary sampling procedures, or bias in terms of samples sent to each laboratory. Two of the laboratories which provided the mycotoxin contamination retrospective data, are private entities and most of their samples were collected and delivered for analysis by clients while samples for the third laboratory were collected by researchers. Heterogeneous distribution of mycotoxin contamination in food and feed materials necessitates employment of appropriately designed and applied procedures for representative sampling [87] and there is no guarantee that the sample were representative since this information was not provided. Further, the purposes for analysis were varied ranging from research, routine monitoring to outbreak of gastrointestinal conditions. It was noted that maize, and total aflatoxin were respectively the most frequently tested food matrix and mycotoxin. All this could introduce a bias in the present study. Consequently, the sample sizes were in most cases inadequate and some important food items and mycotoxins were left out in the present study. Also, the analytical methodology employed by the laboratories were mostly enzyme immunoassays with different limits of detection and quantification, sensitivity and specificity, and were not necessarily those recommended by the food contaminants regulatory organizations. Nevertheless, our data provide a credible evidence of a possible scenario for exposure to dietary mycotoxin hazards.

Lastly, we noticed some areas in the national mycotoxin regulation standards that require some revision. The European standard is quite strict with very low legal limits. For example the maximum limit for aflatoxin M1 in processed dairy products by Kenyan [21,32,37,39] American [50] and FAO/ WHO’s/ *Codex Alimentarius* [88] is 10-fold (500 ppt) compared to the European standard of 50 ppt [52]. The latter further set the limit to 25 ppt for the toxin in infant formulae and milk all this adequately protecting infants and young children. We further observed that many food and feed items consumed locally are not covered by the safety standards. International standards, mostly *Codex Alimentarius* are adopted in their original version without tailoring them for local scenario. Further, although there is high likelihood of some mycotoxin hazards such as ochratoxin A, zearalenone and T-2 toxin, we found no mention of their legal limits set in the national regulatory standards. These agree with Matumba *et al*. [89] and Trench *et al*. [14] who observed that, although various mycotoxins are found in developing countries, more emphasis is put on aflatoxins. In addition, enforcement of mycotoxin regulatory standards is rare in developing countries [23,90]. Protection of population from dietary contaminants, is a function of both establishment of sound regulatory limits and their effective enforcement, and whose driving force lies in a balance between health benefits and food security concerns. In Sub-Saharan Africa, where there is rampant food scarcity, the desire to feed increasing populations erroneously outweighs the health benefits of mycotoxin regulation. High frequency of mycotoxin contamination in Sub-Saharan Africa should be enough incentive for more strict regulation in the region. Nevertheless, recent proposal to review aflatoxin limits by KEBS at the East African Community platform is a welcome move.

## Conclusions

We conclude that animals and humans, including infants, young children and expectant mothers, in Kenya are exposed to an array of dangerous dietary mycotoxin hazards which could lead to serious economic and health implications including cancer. Active surveillance of all dietary mycotoxin hazards observed in this study should be enhanced employing representative sampling plans. More often, concurrent contamination by more than one mycotoxin occurs including masked mycotoxins whose data were not included in the present study. Regulation and future research should therefore focus on multi-mycotoxin analysis techniques, collection of data on toxicological effects of concurrent mycotoxin contamination and consumption pattern, and regulatory limits accordingly set and compliance enforced to protect vulnerable demographic groups such as paediatric, geriatric and sick members of the society.

## Acknowledgements

The Director General (KALRO) and Deputy Director General (Livestock Research) granted permission to publish these data. We thank the mycotoxin-testing laboratories, KALRO-Food Crop Research Institute-Kitale, Bora Biotech Ltd., and Polucon Services (K) Ltd for kindly providing the data used in this study. Mr. Phochunatus Sifuna (KALRO), Mr. George Kimani (Bora Biotech Ltd) and Mr. Charles Maina (Polucon Services (K) Ltd, compiled these data. Mr. David Kinoti and M/s Joana Auma (KALRO) kindly provided statistical and reference management programs respectively. Mr. Nicholus Ndiwa (ILRI) offered training on use of SPSS statistical program. A former science teacher, Mr. R. S. I. Karuku is highly acknowledged posthumously for inspiring the first author to the world of food poisoning, which is the main drive in this communication.

## Supporting information

**S1 Dataset.** Dietary mycotoxin hazards (Excel doc).

## Author Contributions

**Conceptualization**: James Kibugu, Delia Grace, Lindahl Johanna

**Data Curation**: James Kibugu

**Formal Analysis**: James Kibugu, Richard Kurgat

**Funding Acquisition**: James Kibugu, Delia Grace, Lindahl Johanna, Leonard Munga

**Investigation**: James Kibugu, Lindahl Johanna, David Mburu, Leonard Munga

**Methodology**: James Kibugu, Delia Grace, Lindahl Johanna

**Project Administration**: James Kibugu

**Resources**: James Kibugu, Fransisca Lusweti, Bernard Mukasa

**Software**: James Kibugu

**Supervision**: James Kibugu, Delia Grace, Lindahl Johanna

**Validation**: James Kibugu, Delia Grace, Lindahl Johanna

**Visualisation**: James Kibugu

**Writing-Original Draft Preparation**: James Kibugu

**Writing-Review & Editing**: James Kibugu, Delia Grace, Lindahl Johanna, Leonard Munga, David Mburu, Richard Kurgat

